# Genome structure predicts modular transcriptome responses to genetic and environmental conditions

**DOI:** 10.1101/517235

**Authors:** Stephanie Mark, Joerg Weiss, Eesha Sharma, Ting Liu, Wei Wang, Julie M. Claycomb, Asher D. Cutter

## Abstract

Understanding the plasticity, robustness, and modularity of transcriptome expression to genetic and environmental conditions is crucial to deciphering how organisms adapt in nature. To test how genome architecture influences transcriptome profiles, we quantified expression responses for distinct temperature-adapted genotypes of the nematode *Caenorhabditis briggsae* when exposed to chronic temperature stresses throughout development. We found that 56% of the 8795 differentially-expressed genes show genotype-specific changes in expression in response to temperature (genotype-by-environment interactions, GxE). Most genotype-specific responses occur under heat stress, indicating that cold versus heat stress responses involve distinct genomic architectures. The 22 co-expression modules that we identified differ in their enrichment of genes with genetic versus environmental versus interaction effects, as well as their genomic spatial distributions, functional attributes, and rates of molecular evolution at the sequence level. Genes in modules enriched for simple effects of either genotype or temperature alone tend to evolve especially rapidly, consistent with disproportionate influence of adaptation or weaker constraint on these subsets of loci. Chromosome scale heterogeneity in nucleotide polymorphism, however, rather than the scale of individual genes, predominate as the source of genetic differences among expression profiles, and natural selection regimes are largely decoupled between coding sequences and non-coding flanking sequences that contain *cis*-regulatory elements. These results illustrate how the form of transcriptome modularity and genome structure contribute to predictable profiles of evolutionary change.

## Introduction

Evolutionary adaptation to varying environmental conditions starts with genetic variability, often with alternate alleles affecting gene regulation and expression. Consequently, understanding the plasticity, robustness, and modularity of transcriptome responses to genetic and environmental conditions is crucial to deciphering how organisms adapt in nature (Ungerer *et al*. 2007). Gene expression represents the most basic level at which phenotypic plasticity to a perturbation can manifest, and therefore underpins the degree of robustness of higher level phenotypes in response to the same perturbation (de Visser *et al*. 2003; Flatt 2005). Because the transcriptome changes in response both to extrinsic factors (e.g. environmental inputs) and to factors that are intrinsic to the organism itself (e.g. genetic background) (Gagneur *et al*. 2013; Grishkevich & Yanai 2013), we must consider both extrinsic and intrinsic contributions in the dynamism of genetic network composition and its genomic architecture. Consequently, it is crucial to determine how much of the genome is expressed differentially in a plastic manner with sensitivity to environmental conditions versus a genetically deterministic manner independently of environmental conditions versus a non-additive combination of both (Grishkevich & Yanai 2013; Knowles *et al*. 2017). Moreover, it remains generally unclear how modular are distinct gene expression responses and what characteristics of the genome predict their composition and molecular evolution. These questions frame some of the key outstanding issues in connecting transcriptome activity to environmental heterogeneity and the molecular evolution of genomes.

Temperature conditions represent a pervasive extrinsic, environmental factor that influences gene expression and can help reveal the relative roles of plasticity versus robustness of transcriptome profiles (Causton *et al*. 2001; Smith *et al*. 2013). If expression plasticity is adaptive, then we expect organisms to modulate their transcriptomes under chronic developmental exposure to heat or cold stress in a coordinated way to maintain fitness. However, homeostasis may break down at environmental extremes and lead to non-adaptive changes in gene expression that simply reflect a ‘broken’ biological system. Pathways associated with the heat shock response are implicated in physiological buffering to acute heat stress (Lindquist & Craig 1988), but chronic sublethal heat stress may not activate this same stress response. By characterizing profiles of transcriptome change to temperature conditions, we can test the robustness of plastic responses to genetic divergence that reflects genomic evolution in the control over gene expression.

Allelic differences can be thought of as a kind of perturbation, a genetic perturbation, that can expose the sensitivity of gene networks in terms of expression changes (Hecker *et al*. 2009). Expression modulated by *cis*-regulatory alleles may minimize adverse pleiotropic effects and, consequently, modest effects of *cis*-regulatory SNPs might only be pronounced when they accrue over long periods of time to give rise to the kinds of expression differences that accumulate between species (Carroll 2008; Stern & Orgogozo 2008; Wittkopp & Kalay 2012). By contrast, changes to *trans*-acting regulators like transcription factors may lead to many downstream pleiotropic consequences. Consequently, large trans-acting effects might make up a substantial fraction of the genetic variability for gene expression differences among individuals within a species and yet rarely contribute to expression differences between species (Wittkopp *et al*. 2004; Smith & Kruglyak 2008; Stern & Orgogozo 2008; Wittkopp *et al*. 2008; Tirosh *et al*. 2009), because most changes that affect fitness are deleterious and eventually get eliminated by natural selection (Keightley & Lynch 2003). The intermediate timescale of adaptive divergence between populations of the same species thus has the potential to expose whether distinct regulatory architecture must be invoked to describe transcriptome changes across the extremes of timescales from polymorphism within a single population to divergence between species.

In this context, extensive transcriptome analysis of the nematode *C. elegans* in response to heat shock and knock-out mutation began with microarrays (Kim *et al*. 2001), with more recent studies using recombinant inbred lines of wild strains to map polymorphic loci that contribute genotype-dependent responses to temperature (Li *et al*. 2006; Grishkevich *et al*. 2012; Snoek *et al*. 2017; Snoek *et al*. 2019). For example, Li et al. (2006) found that among 496 detectable expression quantitative trait loci (eQTL), trans-eQTL were nearly 8-times as likely as *cis*-eQTL to show genotype-by-temperature responses, with subsequent study reinforcing this pattern (Snoek *et al*. 2017). Moreover, eQTL are found disproportionately on SNP-dense chromosome arms in *C. elegans* (Rockman *et al*. 2010). Grishkevich et al. (2012) reported that constitutively-expressed genes in *C. elegans* tend to have short intergenic regions, consistent with simple *cis*-regulatory controls, and that genes with genotype-dependent expression or genotype-by-environment interactions have longer intergenic regions, consistent with complex *cis*-regulation and a larger mutational target. It remains unknown whether natural selection might be important in shaping genetic variation in these features of *C. elegans*, and whether these properties are general across species.

Here we quantified transcriptome expression for *C. briggsae* nematodes from populations with distinct genetic backgrounds adapted to temperature differences associated with their origins in Tropical versus Temperate latitudes (Prasad *et al*. 2011; Stegeman *et al*. 2013; Poullet *et al*. 2015). Global collections and population genomic analyses of *C. briggsae* wild isolates from Tropical and Temperate regions show that they form distinct phylogeographic groups (Cutter *et al*. 2006; Jovelin & Cutter 2011; Felix *et al*. 2013; Thomas *et al*. 2015). Given this ecological context, along with resources like recombinant inbred line (RIL) libraries and chromosome-scale genome assembly (Ross *et al*. 2011; Stegeman *et al*. 2019), *C. briggsae* represents a valuable system to understand the links between temperature and genetic background in differential gene expression. The exemplar Tropical and Temperate genotypes used as RIL parents, the focus of the present study, exhibit diverse temperature-dependent phenotypic differences consistent with adaptive differentiation of the phylogeographic groups overall, including for fecundity (~2-fold difference at 14°C, ~4-fold difference at 30°C), motility, and gamete development traits (Prasad *et al*. 2011; Stegeman *et al*. 2013; Poullet *et al*. 2015; Stegeman *et al*. 2019). By rearing these animals at hot and cold sublethal temperatures near their fertile limits, as well as under benign thermal conditions, we characterize genotypic and environmentally-induced differential gene expression across the genome. We then describe transcriptome complexity in terms of co-expression modularity to reflect transcriptome plasticity and robustness to environmental and genetic context, demonstrating distinctive genomic spatial distributions, functional attributes, and rates of molecular evolution at the sequence level.

## Materials and Methods

### Experimental design and sequencing

To quantify the genome-wide effects of rearing temperature and genetic background on gene expression, we isolated and sequenced mRNA transcriptomes from *C. briggsae* young adult hermaphrodites of two isogenic strains (AF16 = “Tropical” strain, HK104 = “Temperate” strain) that were reared under “chronic” exposure to 14°C (~150h), 20°C (~65h), and 30°C (~48h) throughout their development from egg to adult. Previous generations of both genotypes had been raised at benign 20°C prior to establishment of eggs for rearing at the treatment temperatures following stage synchronization via standard *Caenorhabditis* sodium hypochlorite (“bleaching”) protocol (Stiernagle 1999), avoiding potential transgenerational effects of stressful temperature on gene expression. After reaching adulthood (checked for young gravid adult hermaphrodites), total RNA was isolated with Trizol extraction and isopropanol precipitation (Tu *et al*. 2015) from mass isogenic cultures of each strain at each rearing temperature with three biological replicates (2 genotypes x 3 rearing temperatures x 3 replications = 18 samples). The mRNA was then separated from small RNA fractions of less than approximately 200 nucleotides using the mirVana kit from Ambion as per the manufacturer’s instructions, and prepared for single-end 100bp sequencing of TruSeq libraries via Illumina HiSeq 2000 (Genome Quebec, Canada) with each of the 18 barcoded samples sequenced across 2 lanes to control for lane effects (Fang & Cui 2011).

We obtained an average of 51.4 million reads per sample (range: 34.5 - 73.4 million) for 925.3 million total reads. Sequences are available in NCBI in project accession PRJNA509247. Over 96% of reads were retained after cleaning and trimming of raw FASTQ files with Trimmomatic 0.36 (894.4 million reads retained), using a seed-mismatch rate of 2, a simple clip threshold of 10, discarding reads <60bp long, and trimming bases from 5’ and 3’ ends if they had phred33 scores lower than 3 (Bolger *et al*. 2014).

### Read mapping and expression counts

For each sample, we mapped reads to the *C. briggsae* genome (WS253) using STAR (Dobin *et al*. 2013), setting the maximum intron size to 5000 bp which includes 99.6% of all intron annotations in the *C. briggsae* reference genome. We applied a liberal mismatch rate of 10 to accommodate potential mapping efficacy differences between the AF16 and HK104 strains due to their genetic differences; the reference genome is based on the AF16 strain, so this liberal parameter choice minimizes the potential for mapping to bias towards the Tropical genotype that could inflate inference of differential expression due to genotype. Over 90% of the 894.4 million total reads mapped to unique locations, in all samples (except one replicate of HK104 at 30°C with 73.86% of 48.4 million reads mapping uniquely), with an average of 45.9 million reads mapping per sample to unique locations (Supplementary Table S1).

We then counted the number of reads that mapped to each exon annotated in the WS253 reference genome with htseq-count (Anders *et al*. 2013) and summed over all exons in a gene to give a raw measure of expression for each gene in each sample. For our analysis, we neglected alternative splicing isoforms, treating them as contributing to expression levels for the same gene, and set the “mode” parameter in htseq-count to “intersection-nonempty” to resolve ambiguity for overlapping genes (Anders *et al*. 2013). Among mapped reads, 82-85% were assigned successfully to a particular gene among the 23,267 genes annotated in the WS253 *C. briggsae* genome in all samples (again excepting one replicate of HK104 at 30°C, with 24.0 million = 58% of reads assigned to genes). Among the reads that were not assigned to genes, most (9% on average) could not be associated with any exon or were counted in multiple locations (8% on average) and less than 0.1% were ambiguous.

### Differential expression analysis

We first visualized gene expression counts in a Multi-Dimensional Scaling (MDS) plot (Nikolayeva & Robinson 2014), which showed strong clustering of most biological replicates within a treatment and differentiation among treatments (Supplementary Figure S1). We then retained only the subset of genes with at least 1 cpm (gene read count per million; using the “cpm” function in edgeR (Robinson *et al*. 2010)) in 3 or more libraries (i.e. in one biological replicate) to exclude 7068 genes with extremely low expression that could bias downstream analysis. It is possible that the genes filtered out at this step might exhibit higher expression at different developmental stages, males, or alternative environmental conditions than those assessed here. To test for statistical evidence of differential expression, we next transformed the expression counts using limma and voom, which performs well in controlling Type I error and in detecting true positives (Smyth 2005; Law *et al*. 2014; Ritchie *et al*. 2015). Preliminary analysis (not shown) found limma to be more conservative than edgeR for our dataset, so we elected to use limma for downstream analysis. Upon applying the voom transformation from the limma package to the remaining set of 16,199 genes, a Q-Q plot showed that the data closely approximated a normal distribution (Supplementary Figure S1).

We then tested these 16,199 genes for differential expression using limma by fitting a linear model to the expression profile for each gene as: expression ~ strain + temperature + strain*temperature interaction. We first tested for significance of the interaction term, and then tested for significance of the main effect terms only if the interaction was non-significant. The model intercept was set as expression for the Tropical strain at 20°C and P-values were adjusted for multiple testing using the Benjamini-Hochberg correction with significance inferred for a false discovery rate (FDR) of 0.05 (Benjamini & Hochberg 1995). To distinguish which genes responded to hot versus cold rearing conditions for genes with a significant effect of temperature (either main effect or interaction effect), we performed post-hoc tests on the individual temperature coefficients (FDR = 0.05). We then classified genes into five mutually exclusive categories based on whether they showed significant differential expression due to genotype (strain) only (“G only” genes), temperature only (“T only” genes), both genotype and temperature as independent main effects (i.e. additive effects; “G&T” genes), a non-additive interaction between genotype and temperature (“GxT” genes), or no differential expression (“no DE” genes).

### Co-expression clustering of gene expression profiles

To capture distinct stereotypical profiles of gene expression differences in response to our temperature and genotype treatments, we performed a co-expression clustering analysis using the Weighted Gene Correlation Network Analysis (WGCNA) package (Langfelder & Horvath 2008). Because WGCNA works best with normally distributed expression values, we again used the voom-transformed expression values for the 16,199 filtered genes. A preliminary hierarchical clustering analysis of the samples rejected batch effects as a source of heterogeneity among samples, instead identifying both genotype and temperature as likely and biologically interesting sources of variation in the data (Supplementary Figure S1). We determined the best soft-thresholding power parameter for our data to be 30 (R^2^ correlation with a scale-free network topology = 0.75) based on fits across a range of values from 1 to 42 (Supplementary Figure S2), which also yielded an acceptable level of mean connectivity (k = 115), which is central to the assumptions of the WGCNA model (Zhang & Horvath 2005).

Running WGCNA yielded 124 initial clusters of genes with similar patterns of expression, which we consolidated further by merging similar modules, defined as those with a correlation of 0.75 or higher with each other (Supplementary Figure S2). This procedure produced 22 co-expression modules plus one pseudo-module (M0) containing the 37 genes that could not be grouped based on expression pattern. The characteristic expression profile of genes in a module is represented by WGCNA as the first principal component in expression space, termed the “module eigengene” (Langfelder & Horvath 2007), which we plotted for each genotype separately as the module eigengene expression values averaged across the three biological replicates as a function of rearing temperature.

We performed statistical overrepresentation tests of Gene Ontology (GO) terms associated with gene lists of each co-expression module using PANTHER (Mi *et al*. 2010), using all four PANTHER lists available for *C. briggsae:* Pathways, GO-slim Molecular Function, GO-slim Biological Process, and GO-slim Cellular Components. P-values were adjusted for multiple testing with the Bonferroni correction.

### Genomic enrichment analysis

*C. briggsae* chromosomes are defined by distinct recombination domains (high recombination arms, low recombination centres, and small tip regions with little detectable recombination), which also correlate with the density of coding genes, repetitive elements and single nucleotide polymorphism (SNPs) (Hillier *et al*. 2007; Ross *et al*. 2011; Thomas *et al*. 2015). We therefore tested whether gene profiles of differential expression or module affiliation were enriched in particular chromosomal regions using Bonferroni-adjusted G-tests, defining arm-center boundaries as in Ross et al. (2011). Analyses of upstream intergenic lengths were log-transformed prior to analysis with ANOVA, excluding genes with overlapping positions in the genome annotation. We used the transcription factor gene designations from (Haerty *et al*. 2008). We also cross-referenced differential expression categories and co-expression module membership with Wormbase-defined *C. elegans* orthologs found have sex-biased differential expression by Ortiz et al. (2014), which we used to test for enrichment with G-tests.

### SNP and molecular evolution analysis

Genotype-dependent differences in expression could result from allelic differences in the local vicinity of genes (*cis*-acting effects; e.g. variants in promoter or nearby enhancer elements) or in distant regulators (trans-acting effects; e.g. variants in the regulation or functional sequence of transcription factors or miRNAs) (Rockman & Kruglyak 2006). The allelic differences contributing to local *cis*-acting regulation are likely to occur in the upstream promoter regions for those genes showing genotype-dependent expression (Grishkevich *et al*. 2012), though there are additional important roles of downstream and intronic regulatory elements in gene expression (Merritt *et al*. 2008). Therefore, we quantified the incidence of single-nucleotide polymorphisms (SNP) between the AF16 and HK104 genomic backgrounds in 500bp upstream and downstream flanking regions of coding sequences, as promoter regions tend to be in close proximity to coding sequences in *Caenorhabditis* (Saito *et al*. 2013).

We called single nucleotide variants between AF16 and HK104 based on Illumina paired-end sequencing of HK104 to ~33x coverage using identical methods of Thomas et al. (2015), yielding 761,531 SNPs and 173,341 indels. Sequences are available in NCBI in project accession PRJNA509247. We calculated the per-bp density of SNPs (π) in the pairwise comparison of AF16 and HK104 in a 500bp window upstream (and downstream) of coding sequences, excluding genes internal to operons (and using just the 5’-most or 3’-most operon gene for upstream or downstream sequence, respectively). 1070 operons comprising 2573 genes were identified based on orthology and synteny with annotated *C. elegans* operons, as in Tu et al. (2015). We also calculated the per-bp incidence of SNPs for different genomic features on a per-gene basis, including non-synonymous sites, synonymous sites, and introns, in addition to the 500bp flanking regions, after masking non-covered and low-quality sites. The effective number of codons (ENC) metric of biased codon usage was calculated for each gene in the *C. briggsae* reference genome WS253 with codonw (J. Peden, http://codonw.sourceforge.net). We used 6911 coding sequence divergence values (dN/dS’) for 1-1 orthologs between *C. briggsae* and *C. nigoni* from Thomas et al. (2015).

## Results

### Widespread genotype- and temperature-dependent differential gene expression

We tested for differential expression across the *C. briggsae* transcriptome in response to three rearing temperature conditions and two genotypes, based on gene expression quantification from 45.7 million uniquely-mapped RNA sequence reads for each triplicate sample on average (824 million total mapped reads; Supplementary Table S1). Over half (54%, n=8795) of *C. briggsae* genes analyzed showed significant differential expression due to genotype, temperature, or both (16,199 genes tested after quality filtering for the genome’s 21,827 annotated coding genes). The majority of these genes had a significant genotype-specific response to temperature (n=4919 “GxT genes”; 56% of 8795 differentially expressed genes; 30.4% of all genes analyzed; Supplementary File S1) (Figure 1A). In contrast to this “complex” GxT pattern, the remaining 3876 differentially expressed genes exhibited a “simple” dependence on genotype, temperature, or additive effects of both (8.8% “G genes”, n=770; 23% “T genes”, n=1987; 13% “G&T genes”, n=1119 genes). Although 64% more genes overall exhibited a simple plastic response to temperature than a deterministic response to genotype (1987 + 1119 = 3106 vs 770 + 1119 = 1889), the abundance of genes with a complex GxT interaction of both factors highlights the important roles of both environmental plasticity and genetic determinism in transcriptome profiles (Figure 1A).

**Figure 1.**
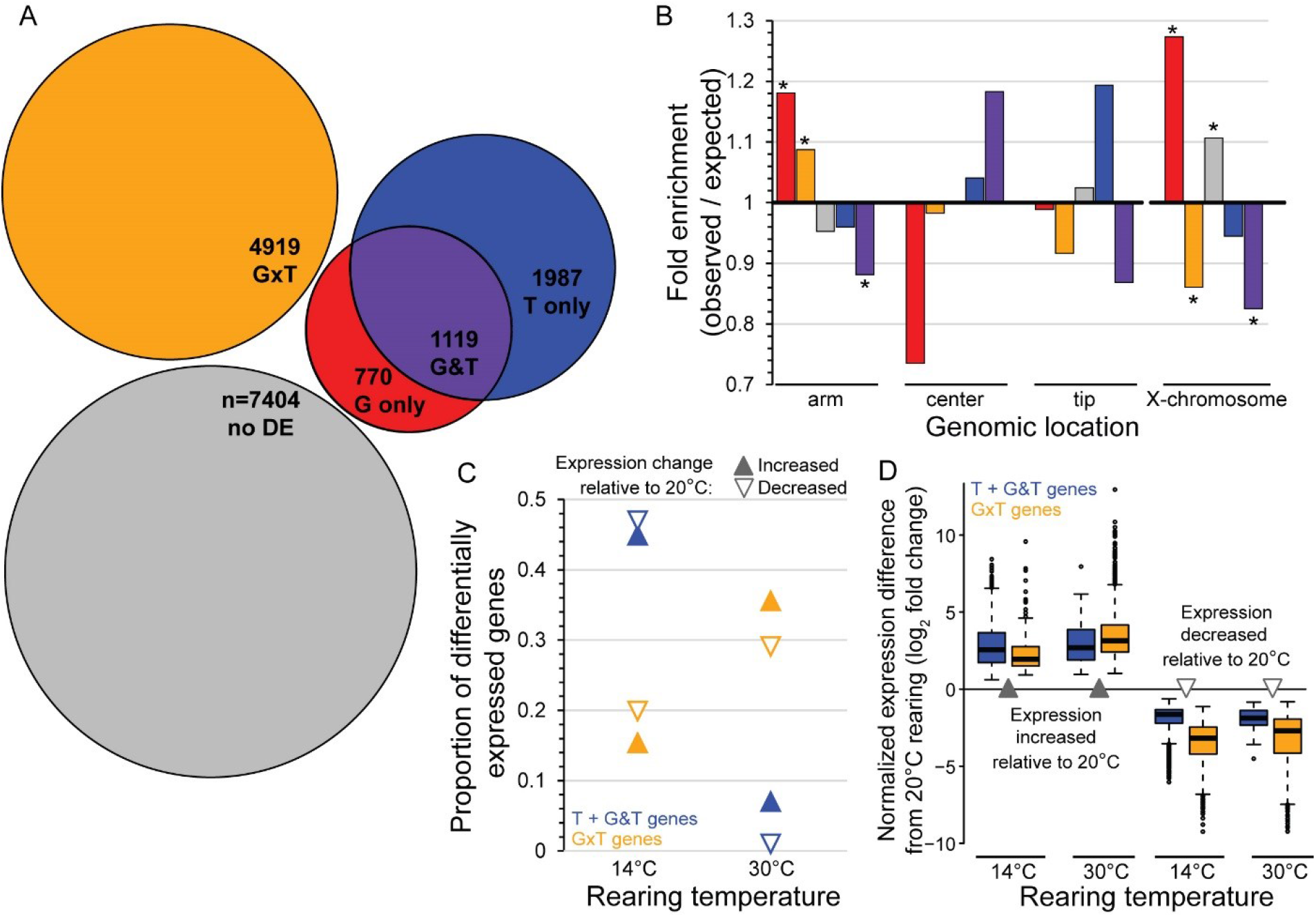
(**A**) Differential expression (DE) analysis identified 54% of 16,199 genes to have significant differential expression (8795 genes at 5% FDR in limma, colored area proportional to gene number). Most DE genes had significant interaction effects (55.9% “GxT”), whereas 12.6% of DE genes had independent additive effects of both genotype and rearing temperature (“G&T”). Other DE genes showed effects of genotype alone or rearing temperature treatments alone (8.8% “G only”; 22.6% “T only”). (**B**) G-only and GxT genes are significantly enriched on autosomal arms, whereas G&T genes are enriched in autosomal centers (colors as in A; * indicates G-test Bonferroni corrected P<0.05). Genes with G-only or no differential expression (“no DE”) are enriched on the X-chromosome, whereas genes with GxT and G&T patterns of differential expression are underrepresented on the X-chromosome (* indicates G-test Bonferroni corrected P<0.05). (**C**) Similar proportions of GxT genes increase vs decrease expression at a given stressful rearing temperature relative to benign 20°C (filled vs empty orange triangles within a temperature condition), but fewer GxT genes show expression differences for cool rearing than for hot rearing (orange triangles for 14°C vs 30°C). By contrast, genes with a non-interacting effect of temperature on expression (T only and G&T genes) show disproportionate response to cool rearing (blue triangles for 14°C vs all other triangles). (**D**) The magnitude of expression change is similar for genes with a non-interacting effect of temperature (T only and G&T genes) under chronic cold stress and chronic heat stress (blue boxes for 14°C vs 30°C; median with interquartile range, whiskers show 1.5x interquartile range). For GxT genes, however, the magnitude of expression increase is greater under heat stress than cold stress (orange boxes for 14°C vs 30°C).

### Distinct genetic responses to chronic heat versus cold stress

Genes with expression influenced by chronic cold stress (14°C) responded differently than genes affected by chronic heat stress (30°C) in terms of the number of genes involved, whether genes increased or decreased expression, and the magnitude of expression change. In particular, cold stress affected expression of 74% of those genes with simple effects of temperature relative to benign conditions at 20°C (2308 of the 3106 T and G&T genes), whereas it was heat stress that altered expression of the plurality of GxT genes (2393 of 4919 genes, 49%) (Figure 1B). Among all these genes that responded to temperature in some way, more genes showed reduced expression at cool temperatures and elevated expression at warm temperatures, compared to benign conditions (Figure 1C; 1.05-fold reduction for T plus G&T and 1.3-fold reduction for GxT at 14°C, 6.8-fold elevated for T plus G&T and 1.2-fold elevated for GxT at 30°C). In terms of the magnitude of differential expression, chronic cold and heat stress were similar for genes with simple expression dynamics (T and G&T genes; Figure 1D; 6.1 to 6.5-fold increase for heat and cold; 3.0 to 3.7-fold decrease for heat and cold). The magnitudes of elevated expression change for GxT genes, however, were much larger under chronic heat stress than under chronic cold stress (hot 8.57-fold vs. cold 3.73-fold increase). Reciprocally, GxT genes that decreased expression under chronic cold stress had a larger magnitude change than under chronic heat stress (hot 6.50-fold vs. cold 9.19-fold decrease). These observations support the idea that distinct genetic networks mediate response to cold versus heat stress, rather than control by a single shared temperature stress response.

### Co-expression modules define gene sets with distinct sensitivities to temperature and genotype

We defined 22 co-expression clusters in the *C. briggsae* transcriptome with WGCNA to capture modules showing distinctive patterns of differential gene expression in response to temperature and genotype differences (Figure 2, Figure 3, Supplementary File S1). The stereotypical expression profile for genes in each co-expression module is represented by its “module eigengene,” defined by the first principal component in expression space (Figure 3). These eigengene profiles illustrate how a given module reflects a dominant trend of genotype-dependence (e.g. M10), temperature-dependence (e.g. M12), additive effects of genotype and temperature (G&T, e.g. M4), or genotype-specific sensitivity to temperature (GxT, e.g. M22) (Figure 3). An average of 46% of genes in a module showed individually-significant differential expression, ranging from a low of just 6% (M13) to a high of 84% (M15) (Figure 3, Supplementary Figure S3). Genes with temperature- and genotype-specific differential expression are concentrated within distinct subsets of modules (Figure 3). Moreover, modules differ in sequence characteristics and in their enrichment with sex-related differential gene expression, as described below.

**Figure 2.**
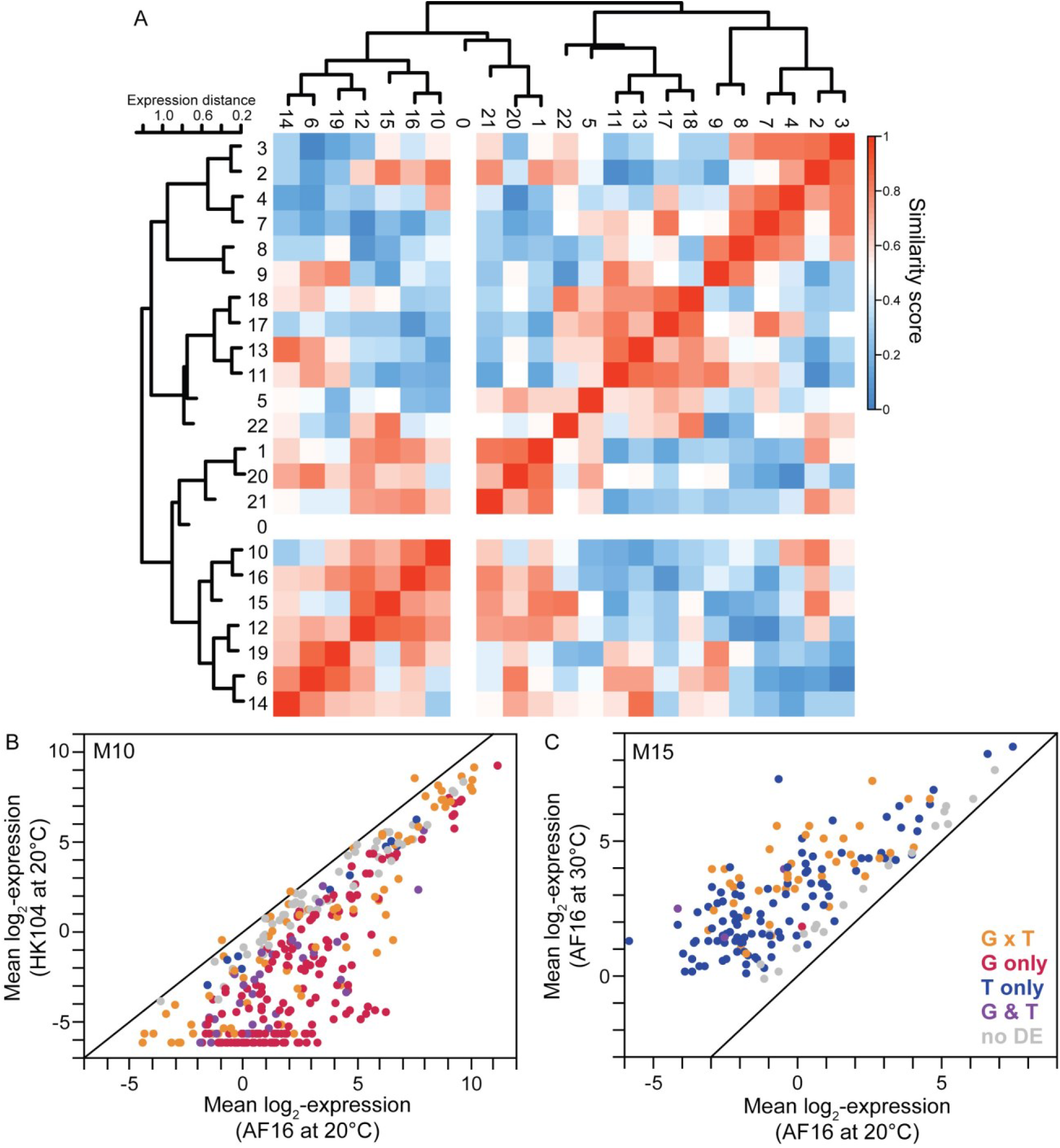
(**A**) WGCNA analysis yielded 23 co-expression modules for 16,199 genes (including module M0 with genes that did not cluster), after merging modules with expression similarity distance <0.25 from an initial set of 124 co-expressed gene sets. The dendrogram and heatmap summarize module expression profile similarity. For example, (**B**) module M10 is comprised of 338 genes with disproportionate representation of genotypic differences in expression, reflected as higher expression by the AF16 (Tropical) genotype. (**C**) Module M15, by contrast, is enriched in genes with individually significant differential expression due to rearing temperature, as reflected in higher expression when reared at 30°C. See also Figure 3.

**Figure 3.**
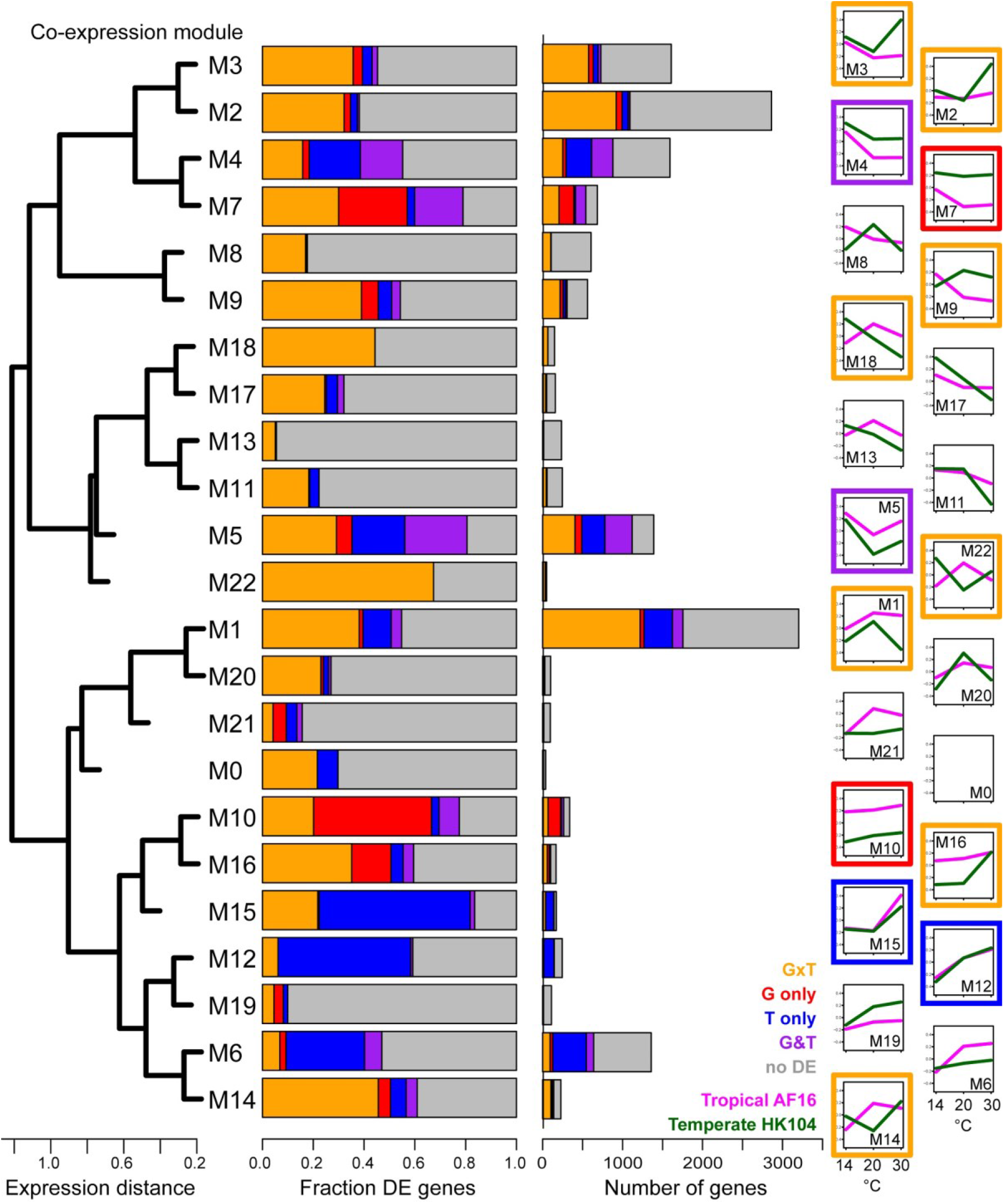
Co-expression clustering of 16,199 genes into 23 modules, reflected in eigengene plots of normalized log2-transformed expression profiles across rearing temperature treatments (14°C, 20°C, 30°C) for each genotype (Tropical AF16, Temperate HK104). Modules range in size from 3203 genes (M1) to 49 genes (M22). Module compositions contained distinctive representation of genes with individually-significant patterns of differential expression (cf. Figure 1); modules with highest incidence of T-only genes indicated with blue (M12, M15), G-only genes in red (M7, M10), G&T genes (M4, M5), and GxT genes with crossing or non-crossing reaction norms (M9, M14, M18, M22; M1, M2, M3, M16) (Supplementary Figure S3).

### Rapid molecular evolution in modules sensitive to genotype or temperature alone

Genotype-dependent expression profiles predominate in just two modules (M7 and M10), which together include 44% (n=342) of all 770 genes with significant ‘genotype-only’ differential expression. Their eigengene expression profiles show limited dynamics across temperatures, with expression for the Temperate HK104 genotype consistently higher than Tropical AF16 in M7 and consistently lower in M10 (Figure 3). M10 is enriched for gene ontology (GO) terms related to extracellular constituents (Supplementary File S2). GO term enrichment in M7 indicates disproportionate representation of genes with nervous system function, including 11 GABA and 11 acetylcholine receptor activity genes, such as the ortholog of *C. elegans* nicotinic acetylcholine receptor *acr-9*. This nervous system enrichment of M7 is salient due to the HK104 and AF16 strains of *C. briggsae* differing in rearing-dependent thermal taxis and locomotion (Stegeman *et al*. 2013; Stegeman *et al*. 2019), a suite of behaviors under neural control.

Genes in module M10 have several other special features compared to other modules: rapidly-evolving protein coding sequences (high dN/dS’), high density of SNPs in replacement sites despite lowest SNP density in introns, the highest enrichment in arm regions of autosomes, enrichment on the X-chromosome, and exceptional rarity in operons (Figure 4, Figure 5A). We observed that genes in M10 also have the least consistent expression among replicates, with very few gene members having orthologs with “oogenic” expression according to Ortiz et al. (2014) (Figure 4A; Figure 5A). These features imply weaker canalization of expression of genes in M10, reflecting either weaker purifying selection or perhaps recent adaptive divergence in average expression levels that has not yet fine-tuned expression variability.

**Figure 4.**
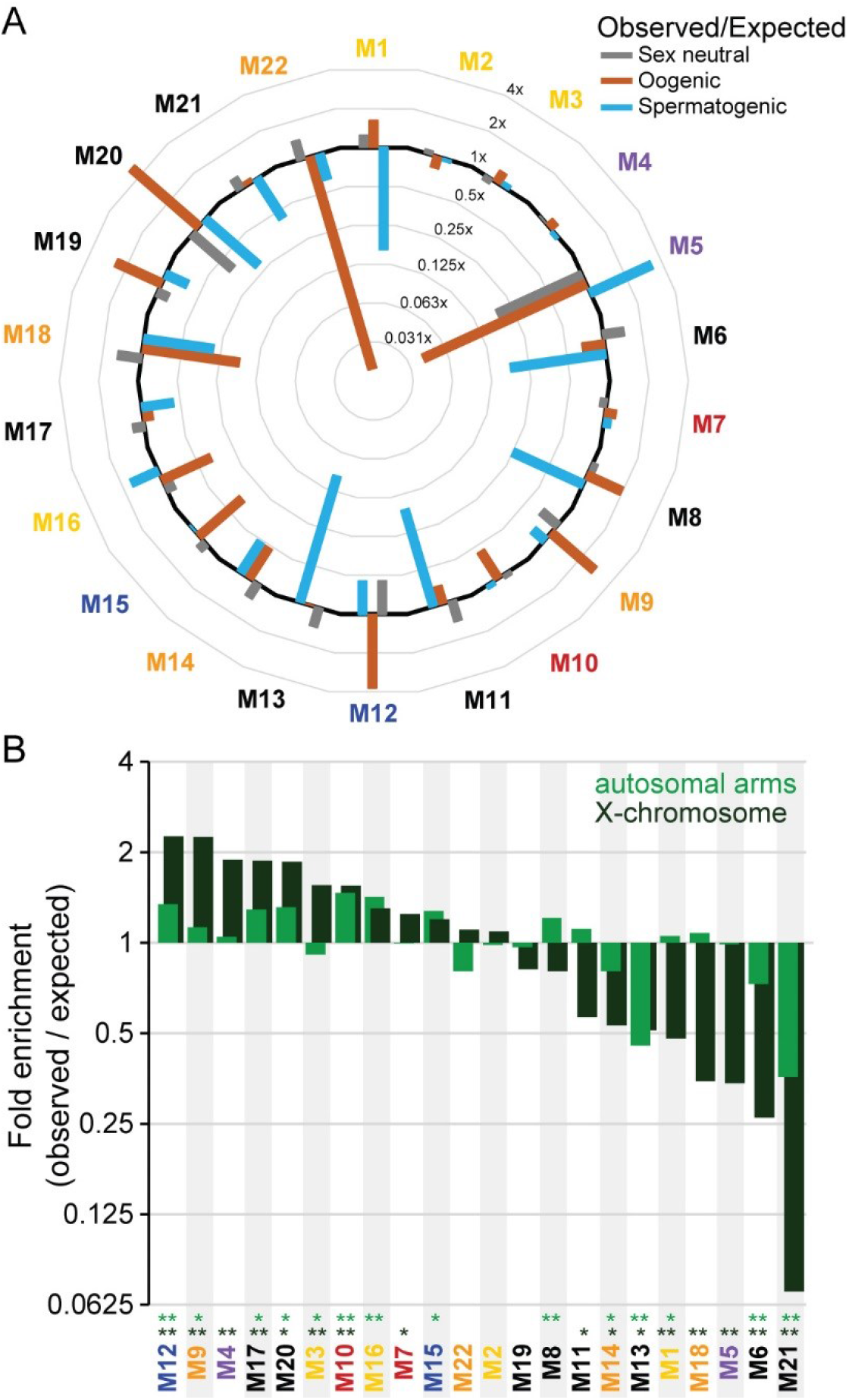
(**A**) Module gene over- and under-enrichment for *C. elegans* orthologs with spermatogenic, oogenic, or sex neutral expression profiles from Ortiz et al. (2014). Log-2 interval scale for observed/expected number of genes per module in radial plot, with black line indicating a value of 1 (outer curve indicates 4-fold enrichment, innermost curve indicates 2^-5^ under-enrichment). (**B**) Module gene over- and under-enrichment across the genome shows biases toward autosomal arms (values < 1 indicate enrichment in autosomal centers) and for X-linkage (values < 1 indicate enrichment on autosomes). Significant enrichment indicated by * (P<0.05 after Benjamini-Hochberg adjustment; FDR = 0.05) and ** (P<0.001 after Benjamini-Hochberg adjustment; FDR = 0.05). Coloring of module names in A and B corresponds to differential enrichment patterns indicated in Figure 3 (blue, T only; red, G only; purple, G&T; orange = GxT with crossing reaction norm profile; yellow, GxT with non-crossing reaction norm; gray, black, differential expression).

**Figure 5.**
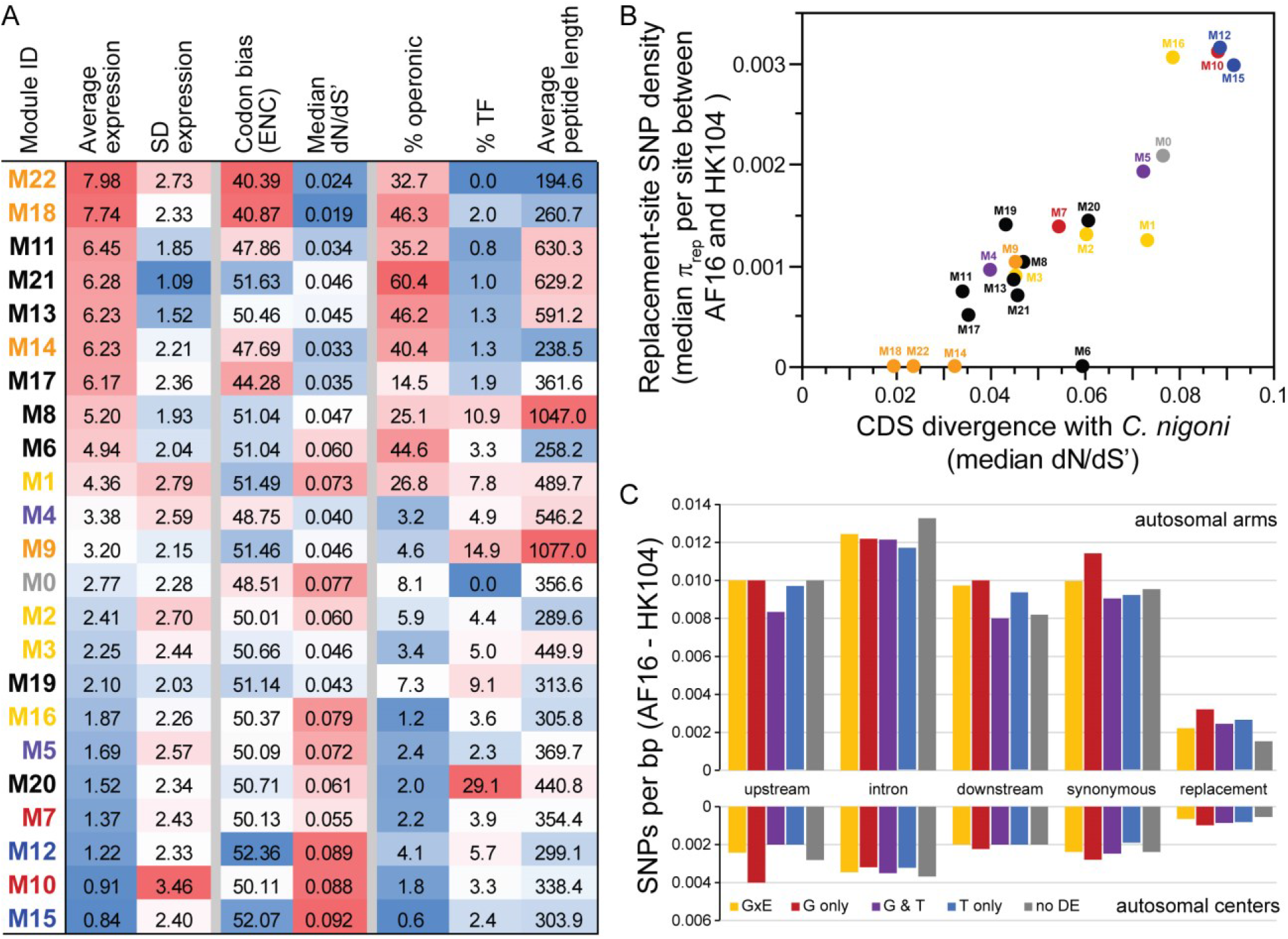
(**A**) Heatmap of module features, sorted by average normalized expression. Modules with high expression profiles tend to contain genes with stronger codon usage bias, greater sequence constraint (low dN/dS’), and more operons. (**B**) Modules with gene orthologs having little coding sequence divergence between *C. briggsae* and *C. nigoni* also have low densities of replacement-site SNPs in coding sequences. Coloring of module names in A and B corresponds to differential enrichment patterns indicated in Figure 3 (blue, T only; red, G only; purple, G&T; orange = GxT with crossing reaction norm profile; yellow, GxT with non-crossing reaction norm; gray, black, differential expression). (**C**) Among genes with distinctive profiles of individually-significant differential expression, linkage to autosomal arms versus centers represents the primary driver of SNP variation with little difference among DE categories for a given genomic site type (1kb upstream of CDS, intronic, 1kb downstream of CDS, synonymous sites).

By contrast to the pronounced genotype-dependent differential expression in modules M7 and M10, two other modules each were comprised of >50% ‘temperature-only genes’ (M12, M15), although they accounted for just 12% (n=247) of the 1987 total T-only gene set (Figure 3). In both modules, eigengene expression is highest at high rearing temperatures across genotypes (Figure 3). In addition, modules M6 and M4 also contained a large fraction of temperature-only genes, and as large modules they also contain a large count of such genes (Figure 3). Modules M12 and M15 have genes with the highest average rates of evolution (dN/dS’) and that occur only rarely in operons (Figure 5). They also have among the lowest average expression levels and codon usage bias (Figure 5A). Module M12 is highly enriched (3.7-fold) for orthologs with an oogenic gene classification in *C. elegans* (Ortiz *et al*. 2014), whereas M15 is depleted of such genes by having 2.6-fold fewer than expected (Figure 4A). Module M12 GO terms show enrichment for genes associated with chromatin, like the ortholog of *C. elegans cec-7*, but with just 8 such genes of the 245 in M12, it is unclear how distinctive a property this is. More enigmatically, M15 shows no GO term enrichment, providing little clue as to whether these heat-sensitive and rapidly-evolving genes act in related functional pathways (Supplementary File S2).

### Sperm gene function associated with both temperature- and genotype-dependence

Two co-expression modules were especially enriched in genes with additive effects of both genotype and temperature (M4, M5; G&T genes), accounting for over half (56%) of all such genes genome-wide (Figure 3). Their eigengene profiles show high expression at low temperatures, with the Tropical AF16 genotype having consistently higher expression than Temperate HK104 in M5 and vice versa for M4 (Figure 3). Interestingly, we found that module M5 is 3.3-fold enriched for orthologs of “spermatogenic” genes from Ortiz et al. (2014), a level unlike any other module (Figure 4A). Genes in M5 also are rare on the X-chromosome and nearly absent from operons, as expected for sperm-related genes (Reinke *et al*. 2000; Reinke & Cutter 2009; Albritton *et al*. 2014), and with fewer transcription factors (TFs) than most modules (Figure 4B; Figure 5A). Moreover, GO term enrichment in M5 indicates a prominent role of genes with phosphatase/kinase activity and glycogen metabolism (Supplementary File S2), including the orthologs of *C. elegans gsp-3/4* and *aagr-1*. Previous expression studies have reported male-biased and sperm-related genes to be enriched for genes with phosphatase/kinase GO terms (Reinke *et al*. 2004; Thomas *et al*. 2012), and some glycoproteins play crucial roles in sperm competitiveness in *C. briggsae* (Yin *et al*. 2018). Sperm fertility is known to show temperature sensitivity differently between the AF16 and HK104 genotypes of *C. briggsae* (Prasad *et al*. 2011; Poullet *et al*. 2015). Thus, the M5 expression pattern implies that universally higher expression for a suite of sperm-related genes, rather than a GxT profile, is associated with the greater sperm fertility at high temperatures observed in the AF16 genetic background.

### Modules enriched for GxE and non-differential expression involved in core biological processes

Eight modules contained an especially large set of GxT genes (M1, M2, M3, M9, M14, M16, M18, M22), indicating a prominent influence of genotype-specific responses to temperature (Figure 3). These eight modules accounted for 71% (n=3477) of all GxT genes genome-wide. The eigengene profiles for four of them show dramatic ‘crossing reaction norms’ such that the Temperate HK104 and Tropical AF16 genotypes exhibit opposite expression responses to rearing temperature (M9, M14, M18, M22; Figure 3). The known genotype-dependence in *C. briggsae* for how sensitive oogenesis is to temperature, with strong reductions of mitotic and meiotic cell counts in the gonad of HK104 animals (Poullet *et al*. 2015), suggests prime candidates among the GxT genes in M9 that has 2.7-fold enrichment for orthologs with oogenic roles (Ortiz *et al*. 2014) (Figure 4A). The other four modules show a much more exaggerated eigengene expression response for just one of the genotypes specifically under high 30°C conditions (M1, M2, M3, M16; Figure 3), rather than crossing reaction norms.

GO terms for core biological processes like mitochondria-related, ribosome-related, and/or translation-related function are enriched in M14, M18 and M22, with M1, M2 and M9 enriched for chromatin and transcription-related GO terms (Supplementary File S2). Genes in modules M18 and M22 also are enriched for orthologs of “sex neutral” genes (Ortiz *et al*. 2014), are enriched in operons, and include few TFs (Figure 4A; Figure 5A). Consistent with these modules involving core biological functions, we also observed the distinctive features of M18 and M22 in having genes with the highest average expression and strongest codon usage bias, while also having the strongest protein sequence conservation (lowest dN/dS’) and the lowest incidence of replacement-site SNPs (Figure 5; Supplementary Figure S5).

The seven remaining co-expression modules consisted primarily of genes that lacked individually significant differential expression, though their module eigengene profiles nevertheless suggest important effects of genetic background and temperature on the stereotypical expression profile (M8, M11, M13, M17, M19, M20, M21). Several of these modules showed GO term enrichment for various metabolic processes (M8, M11, M17, M21) and transcriptional or translational functions (M8, M11, M13, M20). Among these modules, M8, M19 and M20 are extremely enriched for orthologs of “oogenic” genes from Ortiz et al. (2014), but include very few operonic genes (Figure 4A; Figure 5A). M20 also has the highest incidence of TFs (29%) among all co-expression modules, has low average expression, and is enriched for genes on autosomal arms and on the X-chromosome (Figure 4, Figure 5A, Supplementary Figure S5). Genome-wide, TFs are more likely to show no differential expression than other kinds of genes (no DE for 54.5% of TFs vs. 45.2% of other genes; G-test *χ*^2^=29.2, P<0.0001). Module M21 is distinctive in having the highest incidence of genes in operons (60.4%), which are extremely rare on autosomal arms and the X-chromosome (Figure 4, Figure 5A). The 96 genes in M21 have extremely consistent expression across replicates, with most showing no individually-significant differential expression due to either temperature or genotype (Figure 3; Figure 5A). In *C. elegans*, these features are typical of genes that are expressed constitutively across development (Cutter *et al*. 2019).

### Genomic position and differential gene expression

We hypothesized that genomic architectural and molecular evolutionary features might lead to local enrichment of genes with genotype-dependent differential-expression. For example, SNP variation is greater in the high recombination arm domains of autosomes in *C. briggsae* (Thomas *et al*. 2015), and the X-chromosome exhibits a variety of distinctive features compared to autosomes (Ross *et al*. 2011; Cutter 2018). Therefore, we tested for non-random distributions of differentially-expressed genes along chromosomes and between chromosomes. We found that autosome arm domains contained 22% more genes with genotype-dependent expression than expected by chance, and also were slightly enriched for GxT genes (1.04-fold; Figure 1B). Chromosome arms of *C. elegans* also have been reported to contain a disproportionate representation of genes with genotype-dependent differential expression (Denver *et al*. 2005; Rockman *et al*. 2010; Grishkevich *et al*. 2012). By contrast, it was center domains that contained 15% more G&T genes than expected (Figure 1B). Temperature-only genes and genes with no differential expression were randomly distributed between arm and center domains (Figure 1B). Among the 22 co-expression modules, we observed 9 modules to have significant enrichment in arms and 5 enriched in center domains of autosomes (Figure 4B). Thus, gene expression profiles are not spatially independent and genome structural features yield predictable patterns of differential expression within and between chromosomes.

We also found the X-chromosome to be enriched for genes with significant differential expression due to genetic background (G-only genes) as well as for genes with no individually-significant differential expression (X under-representation for G&T and GxT genes; Figure 1B). X-linked biases also held true for co-expression modules (X enriched for 8 modules, autosomes enriched for 8 modules; Figure 4B). Genes from module M21, in particular, are virtually absent from the X-chromosome (Figure 4B), likely associated with the prevalence of operon genes in this co-expression module that also tend to be exceptionally rare on the X-chromosome (Blumenthal *et al*. 2002; Reinke & Cutter 2009). Chromosomes II and IV were distinctive in having no module with significant enrichment or under-enrichment of genes (Supplementary Figure S4). Other autosomes, however, were especially enriched (or under-enriched) for genes in particular modules, for example, genes from M21 were 2.3-fold enriched on Chromosome III and genes from M16 were 2.0-fold enriched on Chromosome V (Supplementary Figure S4). Given the extreme enrichment on Chromosome V for M16 and its genotype-specific expression response at 30°C (Figure 3), it is notable that a quantitative trait locus (QTL) mapping study in *C. briggsae* found QTL on Chromosome V to control differences in heat-sensitive movement behaviors (Stegeman *et al*. 2019).

In *C. elegans*, loci with genotype-dependent expression tend to have longer upstream intergenic regions, interpreted as being consistent with more complex *cis*-regulation of these genes (Grishkevich *et al*. 2012). We observed a similar pattern in *C. briggsae*, with median upstream length of 1367bp for G-only genes versus 1074bp for T-only genes (ANOVA *F*_4,15414_=5.84, P<0.0001, Tukey post-hoc tests on log-transformed upstream intergenic length show G-only > T-only). After partitioning the genomic locations of differentially-expressed genes to account for their non-random distributions in the genome, however, we found that only those G-only genes in autosomal centers have significantly longer upstream intergenic regions compared to T-only genes (arms ANOVA F_4,5653_=0.10, P=0.98; centers F_4,6410_=5.50, P=0.0002, Tukey post-hoc tests on log-transformed upstream intergenic length show G-only > T-only). However, genes in autosomal centers with no differential expression also had longer upstream intergenic lengths than T-only genes and were not significantly different in length to GxT genes or G&T genes. We also find significant variation among co-expression modules in upstream intergenic length (arm ANOVA F_22,5635_=13.59, P<0.0001; center ANOVA F_22,6392_=15.48, P<0.0001), but observe no clear trend between length and the relative composition of genotype- or temperature-dependent genes. Thus, our analysis of *C. briggsae* upstream length distributions does not strongly support the notion that loci with genotype-dependent differential expression have more complex *cis*-regulatory controls.

### Genome structure drives SNP associations with differential expression

We quantified the incidence of single-nucleotide polymorphisms (SNP) for the 761,531 SNPs between the AF16 and HK104 genomic backgrounds in 500bp upstream and downstream flanking regions of coding sequences, as promoter regions tend to be in close proximity to coding sequences in *Caenorhabditis* (Saito *et al*. 2013). We found zero upstream SNPs for 26.2% of the 16,167 genes that had expression and genomic coverage in both AF16 and HK104 (23.0% G-only, 30.9% G+T, 25.6% GxT). Such genes should have no role for *cis*-acting SNPs, suggesting this value as a lower-bound estimate for the incidence of entirely trans-acting regulatory differences that may alter genotype-dependent expression. Moreover, of differentially-expressed genes affected by genotype, 32.6% have zero downstream SNPs, 20.3% have zero intronic SNPs, and 18.6% have zero SNPs in the coding sequence, also consistent with a major role of trans-regulatory control being responsible for the genotype-dependence of differential expression. Consistent with this idea, *C. elegans* shows a predominant role of trans-regulatory control in genotype-dependent differential expression to acute heat stress (Snoek *et al*. 2017).

We further predicted that an important role of *cis*-acting SNPs would be most evident by their enrichment in association with G-only genes (as well as G&T genes and GxT genes), whereas SNPs would be underrepresented in genes with no differential expression or T-only profiles. Genome-wide, we did observe significant differences among differential expression categories in the incidence of SNPs in upstream (ANOVA *F*_4,16141_=7.63, P<0.0001), downstream (*F*_4,16141_=4.75, P=0.0008), and intronic portions of coding genes (*F*_4,16141_=7.82, P<0.0001). Overall, G-only genes have significantly higher SNP densities than other expression classes at replacement sites, synonymous sites, introns and flanking sequences and genes with a GxT pattern of differential expression had a greater density of SNPs than T-only genes only in introns. These results mirror the report by Grishkevich et al. (2012) for *C. elegans* that SNPs are enriched in promoters of genes with genotype-dependent differential expression.

Our findings therefore superficially support the idea of a key role for *cis*-acting SNPs controlling genotype-dependent differential expression. However, we observed that this trend is driven primarily by the enrichment of G-only genes in chromosome arms (Figure 1B), where SNPs are disproportionately abundant for both functionally-constrained and unconstrained sites (Thomas *et al*. 2015). When we account for genomic region, SNP density remains elevated for G-only genes among genes in autosomal centers but not in arms (ANOVA F_4,6729_=3.60, P=0.0062, G-only > other gene classes with Tukey HSD post-hoc test; Figure 5C). Thus, genome structure is an important determinant of inferences about *cis*-acting regulators of genotype-dependent differential expression. We hypothesize that the SNPs in upstream regions of genes in the “SNP deserts” of chromosome centers are more likely to represent causal regulatory variants that modulate gene expression.

### Molecular evolution is decoupled between coding and regulatory sequence regions

Replacement-site SNPs are rarest in the coding sequences of non-differentially expressed genes (in both chromosome arms and centers), consistent with these genes having strongest selective constraint that most effectively eliminates new mutations (Figure 5C). Weaker selective constraint among genes with genotype-dependent differential expression that allows mutations to accumulate could result in their excess of coding SNPs. However, replacement-site divergence between species, which reflects a longer timescale of evolution, is no different between G-only genes and non-differentially expressed genes (median G-only dN/dS’ = 0.0580, no DE dN/dS’ = 0.0511; no significant difference from Tukey’s post-hoc test on log-transformed values). These contrasting patterns for the scale of divergence between phylogeographic groups and between species suggest that relaxed selection on G-only genes might be evolutionarily recent or that adaptive divergence between Temperate and Tropical phylogeographic groups of *C. briggsae* contribute disproportionately to loci with genotype-dependent differential expression.

Consistent with the idea that genes and modules with many SNPs are subject to weaker selective constraints, co-expression modules with high average coding SNP density have low average expression and weak codon usage bias (Figure 5A; Figure 5B; Supplementary Figure S5). Associations were weaker for non-coding flanking regions (Supplementary Figure S5). Modules M10, M16, M12 and M15 were most enriched for coding SNPs (Figure 5B), also exhibiting among the lowest average expression and codon bias. Modules with high coding SNP density also show high long-term molecular evolutionary divergence between species (dN/dS’; Figure 5B), further implicating their constituent genes being subject to weaker selective constraints, or potentially, a greater incidence of adaptive divergence.

Finally, we tested for molecular evolutionary correspondence between coding and non-coding sequence. First, we found that SNPs and interspecies divergence correlate positively across genes for replacement sites, consistent with concordant pressures of purifying selection at both short and long evolutionary timescales on coding sequences (log-transformed π_nonsynonymous_ and dN/dS’, F_1,5741_ = 646.7, P<0.0001). However, interspecies divergence in coding sequence did not correlate with SNP density in non-coding sequences (Supplementary Figure S6). When we analyzed average values for co-expression modules instead of per-gene values, however, we observe positive correlations of non-coding SNP density with both coding SNPs and interspecies divergence (Supplementary Figure S6), suggesting that the distinct gene contents and genomic locations of genes among modules partly contributes to the coding-noncoding correspondence at the module level. Overall, these observations support the idea that selection pressures are largely decoupled between coding sequences and non-coding flanking sequences that contain regulatory elements.

### Muted differential expression role among heat shock proteins

We hypothesized that if heat shock proteins (hsp) modulate transcriptomic responses to chronic temperature stress then we would detect disproportionate differential expression for hsp genes. Of the 24 hsp genes in our expression dataset, only 8 showed significant differential expression, which is less than the genome overall (33% vs. 54%; G-test *χ*^2^ = 4.267, df = 1, P = 0.039), and similar to the genome in differential expression categories (Fisher Exact Test, P = 0.35). This suggests that hsp genes may play a lesser role in temperature stress experienced chronically across development, despite their profound importance to maintaining homeostasis in the face of acute heat stress (Lindquist & Craig 1988); even with acute heat shock, however, few genes show consistent upregulation in *C. elegans* (GuhaThakurta *et al*. 2002).

## Discussion

The *C. briggsae* transcriptome shows widespread differential expression arising from distinct chronic temperatures over development and from distinct genotypes representative of phylogeographic groups from Tropical versus Temperate parts of the world, altering expression for over half of its genes. Genotype-specific responses to temperature represent the most common kind of differential expression (i.e. non-additive genotype-by-environment interactions), with less than a quarter of differentially-expressed genes being sensitive to temperature alone or genotype alone. Our temperature and genotype conditions cluster transcriptome responses into 22 co-expression modules, each comprised of genes with distinctive functional and evolutionary properties that reveal an important role for genome structure in transcriptomic patterns of differential expression.

### The influence of genome structure in differential expression profiles

We found that genome structure plays an important role in shaping the landscape of differential expression of the *C. briggsae* transcriptome, and in the molecular evolutionary features of the corresponding genes. Transcriptome profiles cluster within and between chromosomes, making them susceptible to cryptic correlations with other non-random genomic features. For example, genes showing genotype-dependent expression were enriched on *C. briggsae* chromosomal arms, genomic regions that also are rich in SNPs and with high rates of recombination (Ross *et al*. 2011; Thomas *et al*. 2015). This pattern is reminiscent of the excess of eQTL and loci with genotype-dependent expression on *C. elegans* chromosome arms (Denver *et al*. 2005; Rockman *et al*. 2010; Grishkevich *et al*. 2012). One possibility is that direct selection drives this pattern, as could occur from either adaptive divergence being more prevalent or from purifying selection being weaker for genes on high recombination arms. Alternatively, it might result as a byproduct of linked selection, known to be potent in the *C. briggsae* genome (Cutter & Choi 2010; Thomas *et al*. 2015), whereby elimination of polymorphisms from low recombination centers simply leads to few loci with the potential to show genotype-dependent differential expression.

The higher recombination rate of arm regions means that natural selection favoring a given allele at one locus will be subject to less interference from selection at other loci in the genome (Hill & Robertson 1966; Comeron *et al*. 2008; Cutter & Payseur 2013). Experiments implicate temperature-related adaptive divergence between *C. briggsae* genotypes from Tropical and Temperate latitudes (Prasad *et al*. 2011). Consequently, gene-specific adaptation to distinct ecological conditions should operate more efficiently for genes on arms, which might yield the enrichment of genotype-dependent expression on arms as well as the more rapid sequence evolution of genes on arms. However, it is difficult to exclude a role of linked selection, as the high self-fertilization in *C. briggsae* leaves a substantial imprint on genomic patterns of variation for both synonymous and non-synonymous polymorphisms (Cutter & Choi 2010; Thomas *et al*. 2015). Moreover, if our observations of genotype-dependent differential expression depend primarily on a small number distant trans-acting upstream regulators that influence many target loci (rather than local *cis*-acting allelic variants for many genes), then the bias toward chromosome arms of differentially-expressed genes might simply be a byproduct of non-random distributions of gene functions encoded across the genome.

### Sequence conservation and regulatory controls

The three co-expression modules with the strongest coding sequence conservation also have the highest make-up of GxT genes, which exhibit crossing reaction norms and very high average expression as well as functional enrichment of core biological processes (M14, M18, M22). This finding of especially strong purifying selection implicates either adaptive plasticity in the expression control of these modules or unusually low robustness of expression levels to perturbation from both genotypic and environmental sources. At the other end of the spectrum, modules displaying the highest average rates of sequence evolution have the lowest overall expression and the most pronounced dependence on genotype alone (M10) or temperature alone (M12 and M15). These findings are consistent with expression level as a key determinant of rates of coding sequence evolution, with faster molecular evolution of weakly expressed genes. Similarly, *C. elegans* genes showing non-interaction differential expression tend to have low expression levels (Grishkevich *et al*. 2012). These results can be explained by weaker purifying selection on low-expression genes, though it remains possible that adaptive change might also play a disproportionate role in their molecular evolution.

Among genes located in chromosome centers, those with genotype-dependent differential expression are enriched for SNPs in upstream non-coding regions, consistent with local *cis*-acting alleles affecting their expression. However, long-term coding sequence divergence correlates poorly with non-coding SNP density across genes, implying that the strength of selection on coding sequence variation may be decoupled from *cis*-regulatory genetic variation (Castillo-Davis *et al*. 2004; Jordan *et al*. 2005; Lemos *et al*. 2005; Liao & Zhang 2006; Tirosh & Barkai 2008) or that regulatory elements are too sparse within flanking DNA to leave a clear selective signature with our approach. Nevertheless, the abundance of loci with zero upstream SNPs suggests that distant trans-regulatory control is a profound source of genetic variation in the differential expression patterns that we quantified, consistent with studies of short evolutionary timescales in other systems (Stern & Orgogozo 2008; Wittkopp & Kalay 2012). For example, eQTL analysis for both *C. elegans* and yeast implicates a stronger role for *trans*-relative to *cis*-regulation of genotype-environment interactions (Li *et al*. 2006; Smith & Kruglyak 2008).

### Gene function in differential expression profiles

We found that module M5 was unique in having a large representation of sperm-related genes among its orthologs to *C. elegans* genes with spermatogenesis-enriched function. It includes overrepresentation of genes with phosophatase/kinase activity and is associated with glycogen metabolism, which previous studies show to be especially important in sperm function (Reinke *et al*. 2004; Thomas *et al*. 2012; Wu *et al*. 2012; Yin *et al*. 2018). Both genotype and temperature were important determinants of expression profiles in M5 (Figure 3), implicating the potential for both adaptive divergence and phenotypic plasticity to influence gene responses. Sperm-dependent fertility appears to be especially sensitive to high temperature, with Tropical and Temperate genotypes of *C. briggsae* differing in sensitivity (Prasad *et al*. 2011; Poullet *et al*. 2015), though module M5 shows additive contributions for genetic and temperature effects. As expected for sperm genes (Reinke *et al*. 2000; Reinke & Cutter 2009; Albritton *et al*. 2014), genes from M5 are especially rare on the X-chromosome and virtually absent from operons.

Opposite to the rarity of operon genes in sperm-enriched module M5, fully 60% of the genes in module M21 occurred in operons and yet less than 17% of them had individually significant differential expression. Overall, genes in operons were much less likely to show significant differential expression than non-operon genes (43% of operon vs. 56% of non-operon genes; Fisher exact text P<0.0001). *C. elegans* operon genes, most of which are conserved in *C. briggsae* (Qian & Zhang 2008), are known to show high expression during growth, as for gonad tissue (Reinke & Cutter 2009) and following growth-arrested states (Zaslaver *et al*. 2011), and generally have non-dynamic expression profiles across ontogeny (Cutter *et al*. 2019). These observations are consistent with operon genes being disproportionately robust to both environmental and genetic perturbation.

### Plasticity versus adaptive divergence in expression profiles

In *C. elegans* and *C. remanei*, plasticity dominates the transcriptome response to temperature stress, at least in terms of acute heat shock (Jovic *et al*. 2017; Sikkink *et al*. 2019). We also found in *C. briggsae* that over 90% of differentially-expressed genes changed at least in part due to temperature, but more commonly due to chronic cool rather than warm conditions. If environment-dependent expression responses reflect adaptive plasticity, then our observations suggest stronger canalization of stereotyped cool-rearing expression responses. While the large number of such differentially-expressed genes does not pinpoint the key determinants of temperature-dependent adaptive divergence, we can nevertheless largely rule out the nearly 9500 genes in the genome that show temperature-only effects or no differential expression. Our analysis of *C. briggsae* finds a stronger signal of genotype-dependent differential gene expression than the *C. remanei* study, perhaps reflecting the longer period of divergence between AF16 and HK104 than between the experimental evolution lines for *C. remanei*, in addition to technical differences between the studies (Sikkink *et al*. 2019). Warm conditions causing pervasive genotype-specific responses in *C. briggsae* might reflect adaptive evolution by the distinct genetic backgrounds from Tropical and Temperate regions (Prasad *et al*. 2011). Phylogenetic comparative analysis of differential expression among genotypes and environments could prove fruitful in deciphering whether shared gene networks across species provide common substrate for adaptive divergence and adaptive plasticity in organismal responses to chronic and acute temperature stress.

## Conclusions

Genome-wide differential gene expression is sensitive to both extrinsic temperature conditions and to intrinsic genomic background in the nematode *C. briggsae*, with co-expression modules defining distinctive functional features, genomic distributions and molecular evolutionary patterns of their constituent genes. Most genotype-specific responses occur under heat stress, indicating that cold versus heat stress responses involve distinct genomic architectures. Co-expression modules associated with reproductive function, and which exhibit strong sensitivity to both temperature and genotype, provide candidates for adaptive divergence between Temperate and Tropical phylogeographic groups of *C. briggsae*. The fastest-evolving protein coding sequences correspond to a predominant influence of temperature alone or genotype alone, and have overall low levels of expression across conditions. However, chromosome-scale patterning of nucleotide differences is a key predictor of SNP content of genes, undermining gene-centric causes and *cis*-regulatory inferences for SNP differences across differential-expression classes of genes. These findings highlight the powerful way that genome structure can influence transcriptome profiles to make them susceptible to cryptic correlations with other non-random genomic features.

## Acknowledgements

We are grateful to Rajarshi Ghosh and Leonid Kruglyak for helping to generate and share HK104 genomic sequence. We also thank two anonymous reviewers for constructive feedback on earlier drafts of the manuscript. This work was supported by funds from the Natural Sciences and Engineering Research Council of Canada and a Canada Research Chair to ADC and to JMC.

## Data accessibility

Data used for analysis is provided in NCBI in project accession PRJNA509247 for transcriptome and genome sequences, with online supplement summary tables to be submitted to Dryad, strains are publicly available from the Caenorhabditis Genetics Center.

## Author contributions

SM, JW, JMC and ADC designed research; SM, JW, ES, TL and WW performed research; JMC and ADC contributed reagents/analytic tools; SM, TL, WW and ADC analyzed data; SM, JMC and ADC wrote and edited the manuscript. All authors read and approved the final manuscript.

## Supplementary Information

**Supplementary File S1.** “SuppFile_voomNormFiltLog2Expr_DE_modules.csv” contains log-2 normalized expression values for each of the 16,199 genes analyzed in each replicate sample, as well as the category of differential expression (G-only, T-only, G&T, GxT, noDE) and name of the co-expression module (M0 through M22). Columns labeled with sample name (treatment-replicate), where treatment is a combination of genotype and rearing temperature for each of three biological replicates (“AF” = Tropical AF16 genotype, “HK” = Temperate HK104 genotype; 14=14°C rearing, 20=20°C rearing, 30=30°C rearing).

**Supplementary File S2.** “SuppTable_GO.xlsx” contains lists of gene ontology term enrichment with summary statistics for different PANTHER GO-slim categories for each co-expression module.

**Supplementary Table S1.** Number and percentage of reads that mapped to unique genomic locations with STAR.

| Sample<br>(treatment-rep)* | Number of uniquely<br>mapped reads | % of reads that<br>uniquely mapped |
| --- | --- | --- |
| AF14-1 | 32007586 | 93.35% |
| AF14-2 | 47012080 | 93.84% |
| AF14-3 | 54523984 | 94.15% |
| AF20-1 | 57787850 | 94.17% |
| AF20-2 | 62122726 | 93.43% |
| AF20-3 | 43596994 | 93.75% |
| AF30-1 | 66295835 | 93.74% |
| AF30-2 | 35618842 | 93.20% |
| AF30-3 | 33087508 | 90.10% |
| HK14-1 | 51264144 | 93.39% |
| HK14-2 | 43616826 | 93.65% |
| HK14-3 | 48035023 | 92.74% |
| HK20-1 | 30262908 | 90.76% |
| HK20-2 | 49528582 | 93.95% |
| HK20-3 | 41332043 | 94.26% |
| HK30-1 | 51537189 | 91.44% |
| HK30-2 | 40748065 | 92.22% |
| HK30-3 | 35632338 | 73.62% |

\*AF=AF16, HK=HK104 genotypes; 14=14°C, 20=20°C, 30=30°C rearing

**Supplementary Figure S1.**
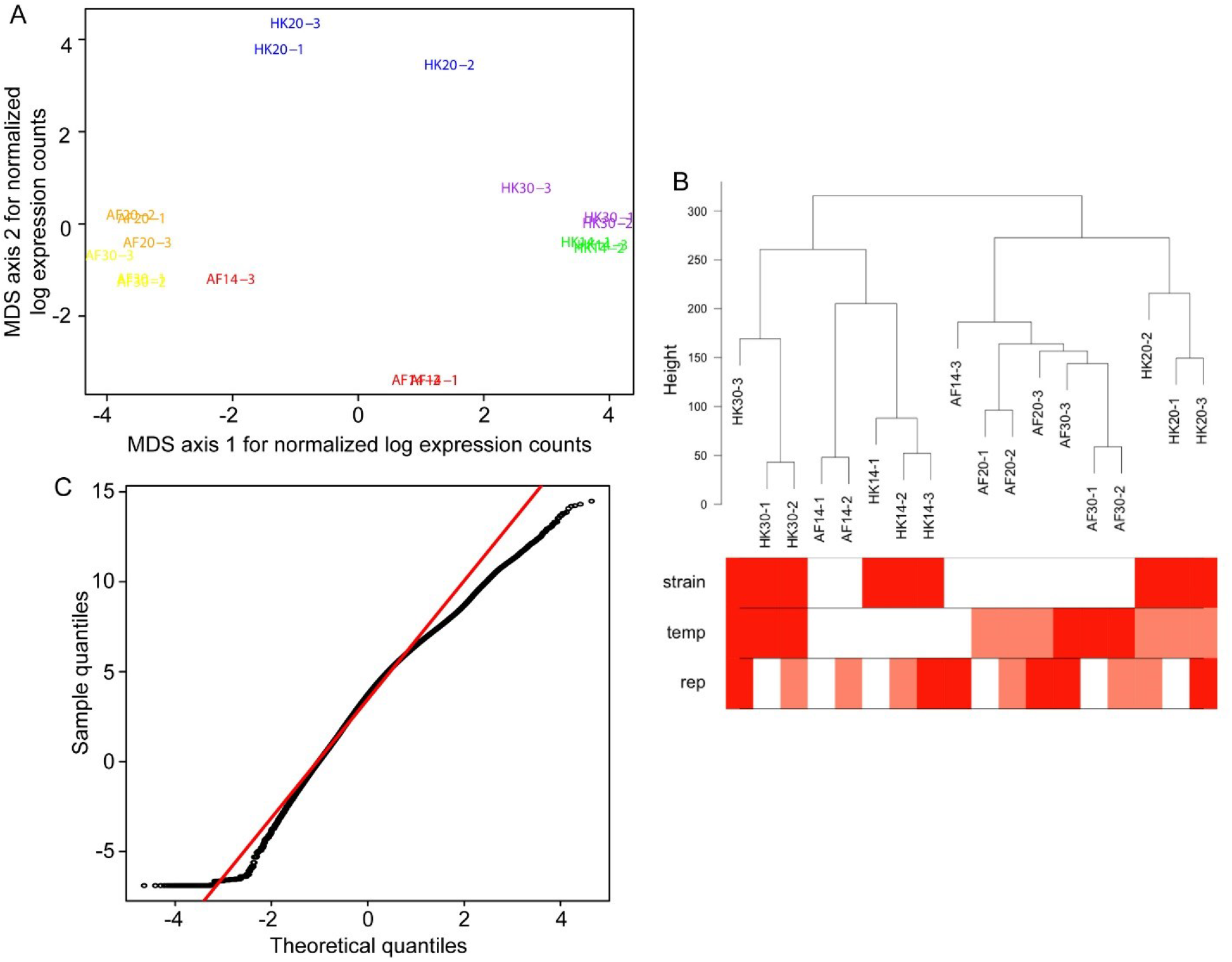
(A) Multi-dimensional scaling plot (MDS) of filtered, normalized, and log-transformed count data for each sample. Sample labels (treatment-replicate) are colored by experimental treatment for a given genotype and rearing temperature combination (“AF” = Tropical AF16 genotype, “HK” = Temperate HK104 genotype; 14=14°C rearing, 20=20°C rearing, 30=30°C rearing). The x- and y-axes show the two principal components that explain most variation. Biological replicates that cluster together in the plot are more similar to each other, indicating consistency across replicates. (B) After filtering out genes with very low to no counts, TMM normalization for different library sizes, and log-transforming count data, a dendrogram reveals similarity of samples within strain and within temperature (“temp”), but not replicate (“rep”); red shading along a row indicates shared strain, temperature or replicate value. This suggests data heterogeneity among samples due to experimental treatments, not batch effects. (C) Quantile-quantile plot for normalized, voom-transformed count data shows good approximation to a normal distribution (red line).

**Supplementary Figure S2.**
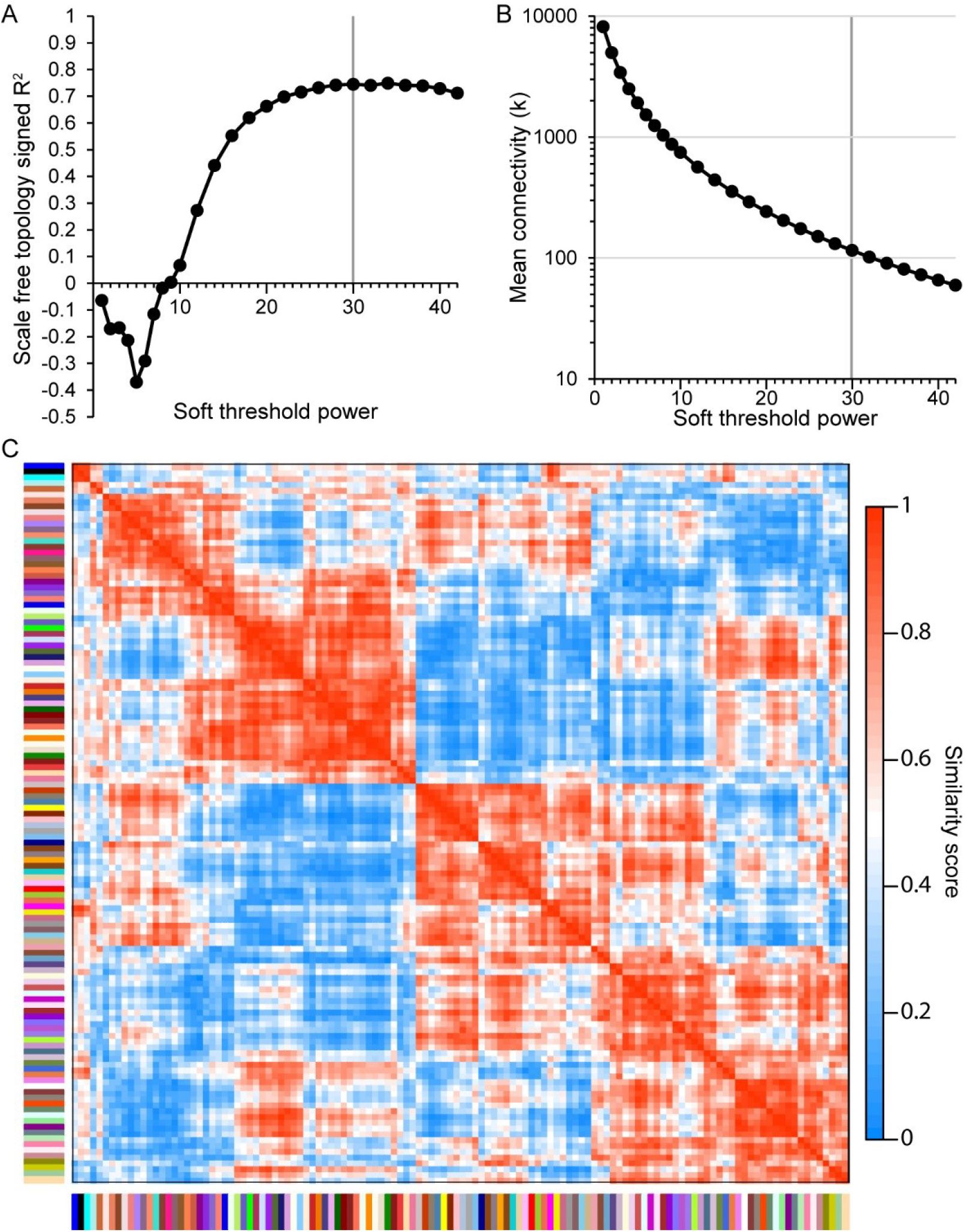
(A) Analysis of soft-thresholding powers revealed 30 to be the power at which the scale-free fit is maximized (R^2^ = 0.75) and most closely approximates a scale-free network. (B) Analysis of a range of soft-thresholding powers revealed a value of 30 to have a mean connectivity of at least 100 (k = 115), while also maximizing fit to a scale-free network. (C) Clustering of 16,199 genes with WGCNA into a preliminary set of 124 co-expression clusters (red in heatmap indicates maximum similarity and blue no similarity; color bars along the x- and y-axes correspond to the 124 clusters). Clusters with similarity distance <0.25 were merged, leading to the 22 co-expression modules (plus pseudo-module M0) analyzed in the main text.

**Supplementary Figure S3.**
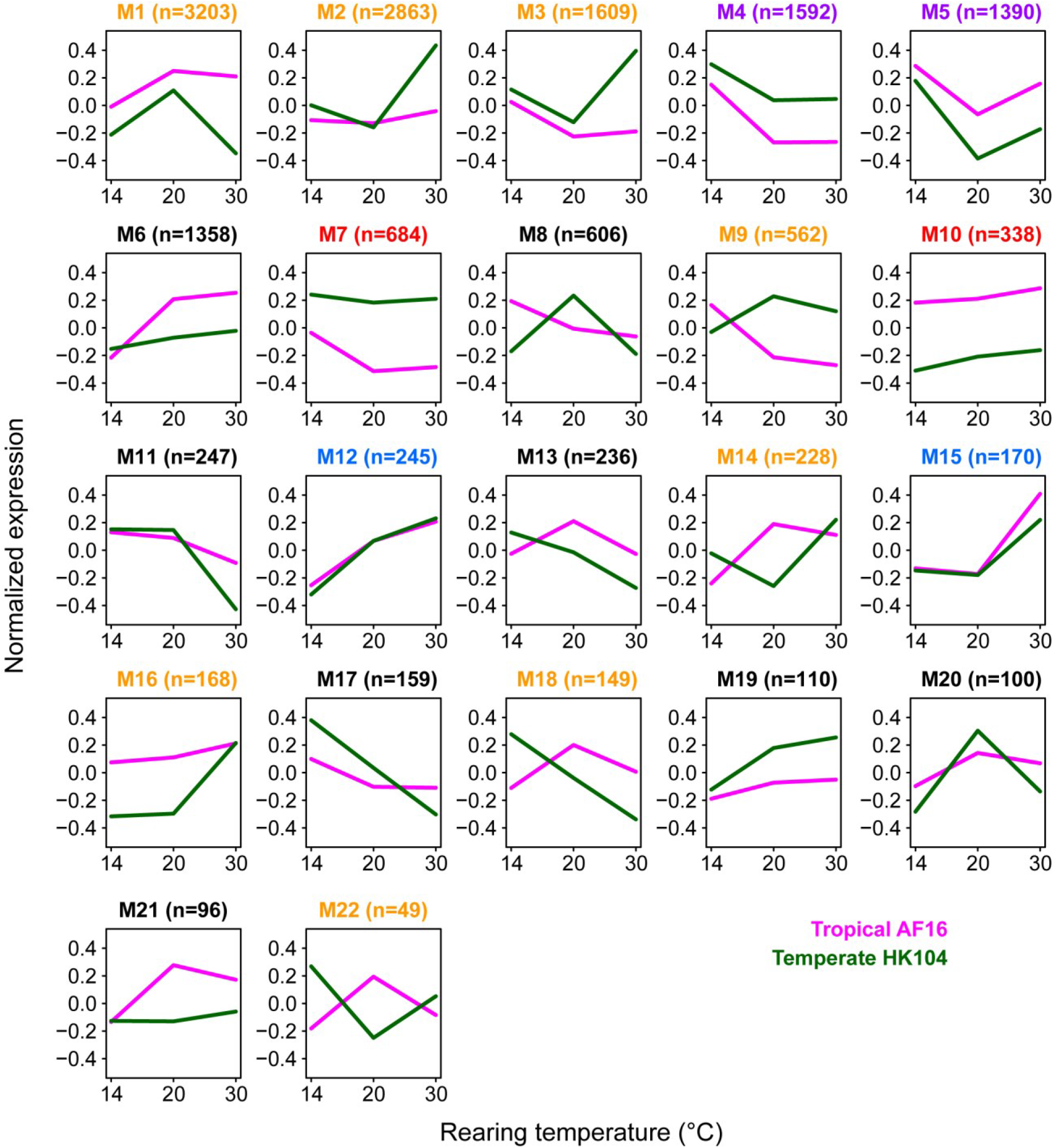
Co-expression module eigengene plots of normalized, log2-transformed expression across temperature treatments for each genotype (pink AF16, green HK104). Module names colored as in Figure 3 of the main text according to representation of individually significant differentially expressed genes within the module.

**Supplementary Figure S4.**
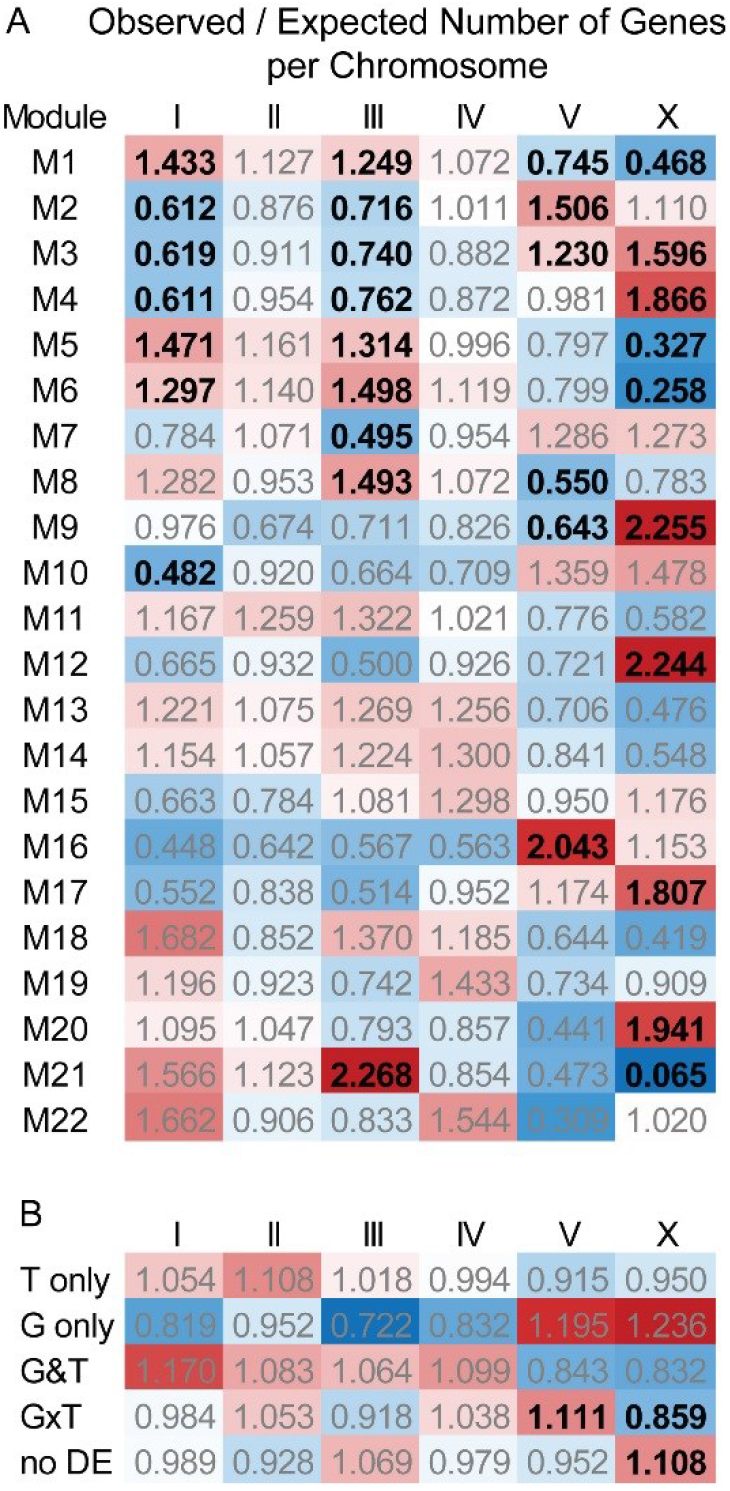
Enrichment (observed/expected) number of genes on each chromosome for modules (A) and differential-expression categories (B). Values in black bold text indicate significant enrichment or under-enrichment after Bonferroni multiple-test correction (*χ*^2^ test df=1 with α=0.05/132 for modules, α=0.05/30 for categories).

**Supplementary Figure S5.**
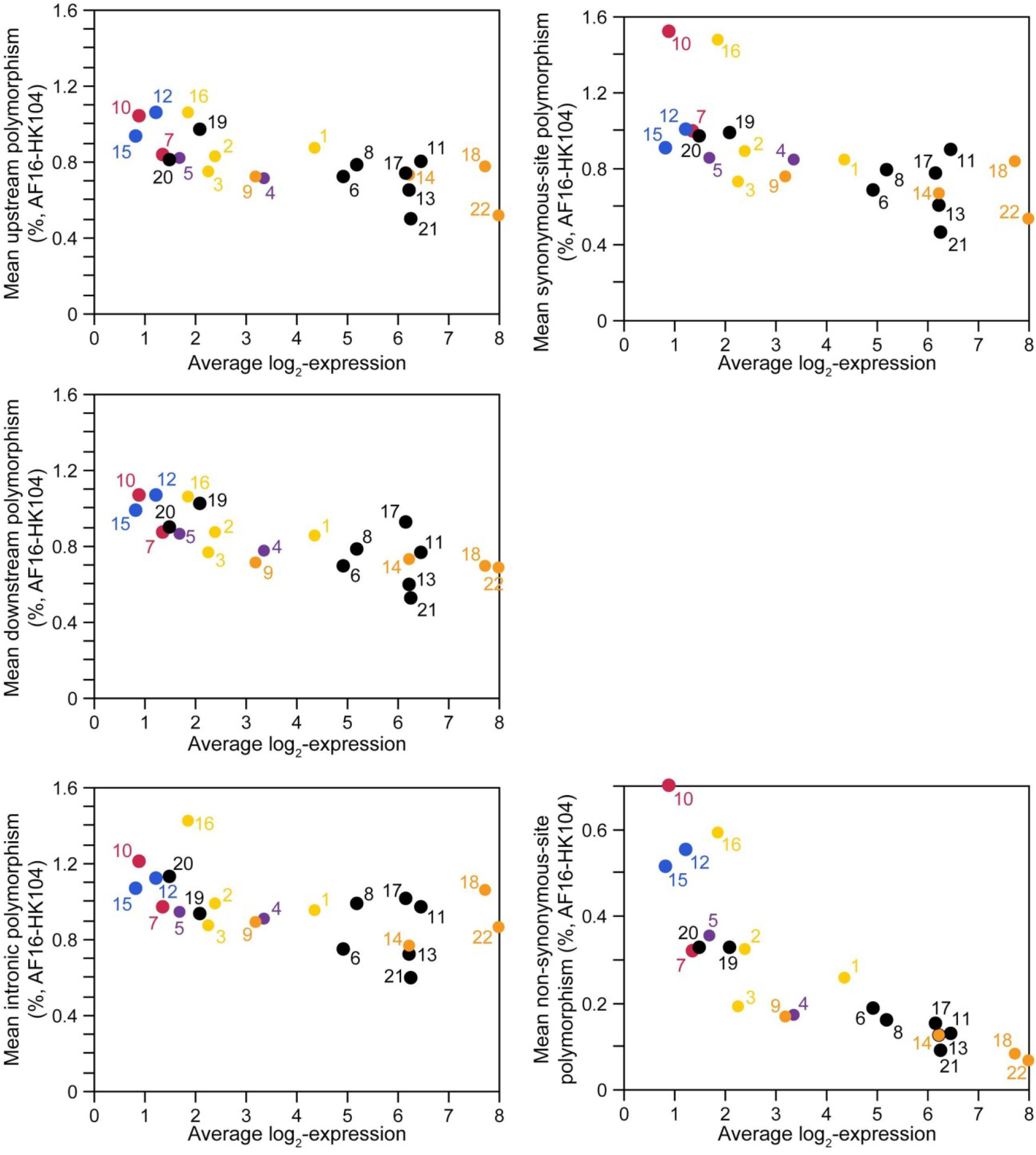
SNP density in upstream, downstream, and intronic non-coding locations of genes and at synonymous and non-synonymous sites within coding sequences, averaged for genes within each co-expression module as a function of average module expression. Correlation across modules: module mean π_nonsynonymous_ × average expression Spearman ρ = −0.92, P<0.0001; π_nonsynonymous_ × ENC ρ = 0.418, P=0.048; module mean π × average expression Spearman ρ, ρ_upstream_ = −0.72, P<0.0001; ρ_downstream_ = −0.77, P<0.0001; ρ_intronic_ = −0.54, P<0.0082. Points are labeled and colored with module name as in Figure 3 of the main text.

**Supplementary Figure S6.**
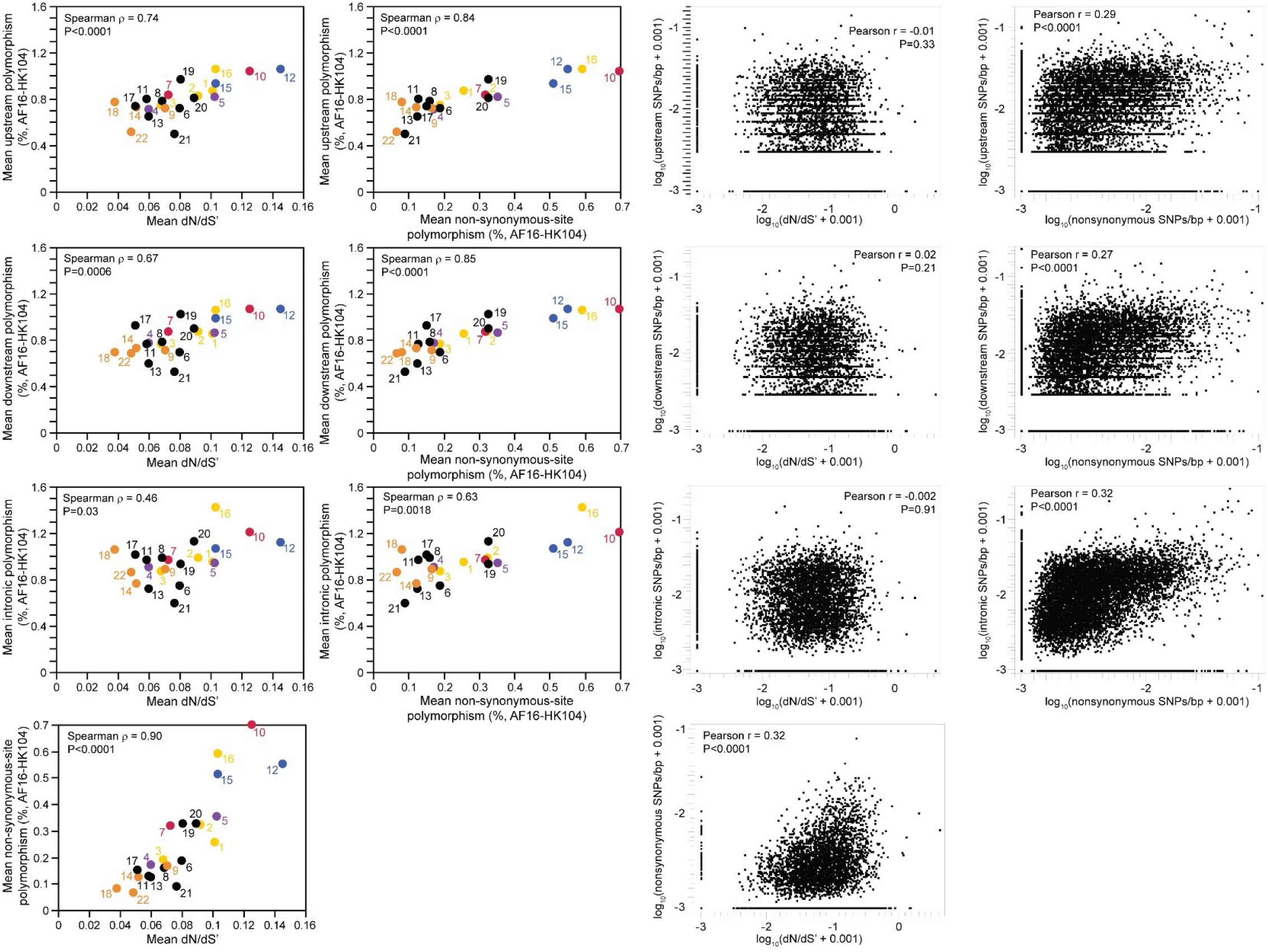
SNP density in upstream, downstream, and intronic non-coding locations of genes and at non-synonymous sites within coding sequences, averaged for genes within each co-expression module (left panels) or per gene (right panels) as a function of average module interspecies divergence (dN/dS’) or non-synonymous site substitution. Modules are labeled and colored as in Figure 3 of the main text.

